# A novel role of VisP and the Wzz system during O-antigen assembly in *Salmonella* Typhimurium pathogenesis

**DOI:** 10.1101/318089

**Authors:** Patrick da Silva, Fernanda Z. Manieri, Carmen M. Herrera, M. Stephen Trent, Cristiano G. Moreira

## Abstract

*Salmonella enterica* serovars are associated with diarrhea and gastroenteritis and are a helpful model for understanding host-pathogen mechanisms. *Salmonella* Typhimurium regulates the distribution of O-antigen (OAg) and presents a trimodal distribution based on Wzy polymerase, Wzz_ST_ (long chain length OAg, L-OAg) and Wzz_fepE_ (very long chain length OAg, VL-OAg) co-polymerases; however, several mechanisms regulating this process remain unclear. Here, we report that LPS modifications modulate the infectious process and that OAg chain length determination plays an essential role during infection. An increase in VL-OAg is dependent on Wzy polymerase, which is promoted by a growth condition resembling the environment of *Salmonella*-containing vacuoles (SCVs). The virulence and stress-related periplasmic protein (VisP) participates in OAg synthesis, as Δ*visP* presents a semirough OAg phenotype. The Δ*visP* mutant has greatly decreased motility and J774 macrophage survival in a colitis model of infection. Interestingly, the phenotype is restored after mutation of the *wzz_ST_* or *wzz_fepE_* gene in a Δ*visP* background. Loss of both the *visP* and *wzz_ST_* genes promotes an imbalance in flagellin secretion. L-OAg may function as a shield against host immune systems in the beginning of an infectious process, and VL-OAg protects bacteria during SCV maturation and facilitates intramacrophage replication. Taken together, these data highlight the roles of OAg length in generating phenotypes during *S.* Typhimurium pathogenesis and show the periplasmic protein VisP as a novel protein in the OAg biosynthesis pathway.

**Author summary:** *Salmonella* modifies its LPS, specifically the O-antigen length, to adapt itself to distinct intestinal environments. These LPS modifications may provide a way for this bacterium to avoid complement activation in the intestinal lumen, improving *Salmonella* pathogenesis. This process is essential for a successful infection, and our investigation into these specific details regarding LPS in this foodborne pathogen will elucidate different aspects of the host-pathogen association.

## Introduction

Bacteria-host interactions are a natural synergistic relationship. However, this interaction may diverge, leading to distinct consequences that range from essential and beneficial cooperation to deadly outcomes. *Salmonella* is a human pathogen that is responsible for intestinal infections and typhoid fever. Salmonellosis is one of the most common and broadly found foodborne illnesses in the world. Yearly estimates include approximately ten million human cases worldwide, resulting in more than 100,000 deaths (1). *Salmonella enterica* serovar Typhimurium is a major foodborne pathogen responsible for gastroenteritis and complications such as serious invasive non-typhoidal *Salmonella* (iNTS) and is frequently reported in Sub-Saharan regions (2, 3). *S.* Typhimurium is acquired through contaminated food ingestion and has mechanisms to survive the low pH milieu and bile salts of the stomach, allowing it to reach the intestine (4). Once in the intestinal lumen, *S.* Typhimurium employs its flagella, a complex multiprotomeric structure, to move toward a nutrient-rich environment, such as the epithelial mucosa, and thus establish a colonization process (5, 6). The goblet cells in the normal intestine accumulate multiple mucous storage vesicles. *S.* Typhimurium can benefit from this propitious environment by inducing inflammation to increase the secretion of mucin and glycoconjugate, high-energy nutrient sources, by the goblet cells (5, 7). The release of glycoconjugates from goblet cells creates an attractant gradient for *S.* Typhimurium, causing *S.* Typhimurium to move in its direction and hence initially attach to epithelial cells (5).

The inflammatory process can be initiated by different *S.* Typhimurium membrane structures, such as flagellins, which are flagella subunits (8-10); type-three secretion system (TTSS) structural proteins (11-16); lipopolysaccharide (LPS) (17, 18); and other effector proteins encoded in *Salmonella* pathogenicity islands (SPI) (19-22). Fimbriae and other adhesins help *S.* Typhimurium initiate contact with epithelial cells by binding to membrane mannosidic glycoproteins, which anchor the bacterium (23, 24).

Bacterium-host cell interactions enable SPI-1 TTSS to inject effector proteins into the epithelial cell, which modify the cell’s structure by ruffling its membrane, culminating in bacterial engulfment (25). *S.* Typhimurium can live and replicate within the intracellular environment by manipulating cellular vesicle trafficking to obtain nutrients (4). Intracellular settling and colonization are classically mediated by SPI-2 TTSS protein effectors (4, 26, 27) and by SPI-3 proteins associated with metabolic adaptations, such as ion transport (28). Cytokines released by epithelial cells into the bloodstream in response to an inflammatory process are responsible for recruiting polymorphonuclear cells (PMN) such as neutrophils to eliminate pathogens (21, 29). Therefore, the epithelial cell invasion route helps *S.* Typhimurium evade an immune response. However, *S.* Typhimurium can overcome macrophage phagocytosis by employing the SPI-2 TTSS machinery, which enables bacterial survival and intracellular replication (4).

The bacterial membrane plays an essential role in all of these pathogenic processes. Its structure provides protection against host defense mechanisms (30, 31); physical stability for multiprotomeric complexes, such as flagella (32, 33), fimbriae (34) and TTSS (35); and bacterial homeostasis by controlling ion transport (36-38) and sensing environment stimuli (39-41). Individual bacterial membranes activate cascade responses to better adapt themselves to different situations (42-51). LPS is the major constituent of the outer leaflet of the outer membrane (OM) of *S.* Typhimurium. LPS is composed of three chemically distinct parts: the membrane-embedded lipid A molecule, which is the hydrophobic and immunogenic portion of LPS that anchors LPS to the OM; the oligosaccharide core; and the highly immunogenic O-antigen (OAg). The OAg molecule is a polysaccharide chain composed of four-sugar repeat units (RU) that externally protrude from the bacterial membrane (52). Gram-negative bacteria synthesize the OAg RU linked to undecaprenyl-phosphate (UndP), an essential bacterial lipid that “carries” various hydrophilic precursors across the membrane, on the cytoplasmic side of the inner membrane. The flippase Wzx then transports UndP-linked OAg RU to the periplasmic face of the IM. The OAg RUs are then used as building blocks for chain elongation, a process mediated by the Wzy polysaccharide polymerase (53-55). Initially, Wzy binds to the UndP-linked RU and then adds another RU or a growing chain of an OAg to the educing end of the OAg RU in a catalytic distributive mechanism (53, 54, 56). In *S.* Typhimurium, the Wzz polysaccharide co-polymerases (PCPs) then serve as a molecular ruler that determines the final length of the OAg chains. *S*. Typhimurium OAg chains follow a trimodal distribution (35) of short (S-OAg, less than 16 RUs), long (L-OAg, 16 to 35 RUs), and very long forms (VL-OAg, more than 100 RUs). The Wzz_ST_ and Wzz_fepE_ proteins regulate L-OAg (57-59) and VL-OAg synthesis, respectively (60).

Recently, our group reported that a novel membrane protein, virulence- and stress-related periplasmic protein (VisP), enables *S.* Typhimurium survival upon encountering stress factors during pathogenesis (61). VisP has an important role in intramacrophage survival and interacts with the lipid A modification enzyme LpxO (61), an Fe^3+^/α-ketoglutarate-dependent dioxygenase (62, 63) related to macrophage evasion (17, 52). VisP was initially identified as a member of the bacterial oligonucleotide/oligosaccharide-binding fold (BOF) family (61) and was predicted to bind oligosaccharides, such as N-acetylglucosamine and N-acetylmuramic acid (64).

In this study, we investigated the role of VisP in the Wzy-dependent pathway of OAg biosynthesis and determined the importance and consequences of the longer forms of OAg chains for *S.* Typhimurium pathogenesis. Furthermore, we also examined how different OAg forms directly affect protection against host defenses during *in vivo* infections and other pathogenic mechanisms, such as flagella-mediated motility.

## Results

### OAg changes are driven by intracellular conditions

*S.* Typhimurium is a facultative intracellular pathogen. LPS is one of a number of known pathogen-associated molecular patterns (PAMPs), and bacteria are recognized by the innate immune system via interactions with these PAMPs. We performed OAg extraction from cells cultured under different growth conditions to observe different luminal or intracellular conditions that trigger OAg chain length changes in *S*. Typhimurium. LB medium was employed as a rich nutrient condition to mimic the intestinal lumen (44, 65). We also harvested bacteria from N-minimal medium under nutrient-depleted conditions, such as low Mg^2+^ (10 μΜ) and phosphate (1 mM KH_2_PO_4_) concentrations, at a mildly acidic pH of 5.0; these conditions are similar to those of the *Salmonella-*containing vacuole (SCV) environment (39, 46, 66-71). The OAg profile of the wild-type (WT) strain presented significant increases in VL- and S-OAg after growth in N-minimal medium compared to the levels after growth in LB (Fig 1A, lanes 2 and 3; see also S1A and S1B Figs). Under both growth conditions, the three distinct modal forms (VL-, L- and S-OAg) were well distributed; however, the VL- and S-bands were the most intense and easily distinguished under N-minimal growth conditions (Fig 1A, lanes 2 and 3; see also S1A and S1B Figs). Apparently, low nutrient conditions, such as the SCV environment, are favorable to more prominent bands of VL-OAg than the lumen conditions of *S.* Typhimurium. Similar VL-OAg chain augmentation was previously described for *S*. Typhimurium C5 growing in inactivated guinea pig sera and iron-limited conditions (31). We measured the expression levels of OAg chain biosynthesis pathway genes to better understand the observed changes in OAg final length. The expression of the *wzy* gene, which encodes the OAg polysaccharide polymerase (53-55, 57, 72), was assessed under both growth conditions for the WT strain, the single mutants Δ*visP*, Δ*wzz_fepE_*, and Δ*wzz_ST_*, and the double mutants Δ*visP*/*wzz_fepE_* and Δ*visP*/*wzz_ST_* (Fig 1B). The OAg profile of the WT strain (Fig 1A, lanes 2 and 3; see also S1A and S1B Figs) matched *wzy* gene expression levels because *wzy* was upregulated in the N-minimal growth condition (Fig 1B). Therefore, *S.* Typhimurium may remodel its OM landscape within SCVs in response to its environment by increasing the VL- and S-OAg forms, according to the observations under growth conditions resembling SCVs (N-minimal medium). Furthermore, this remodeling may be mediated by Wzy polymerase overexpression.

**Fig 1.**
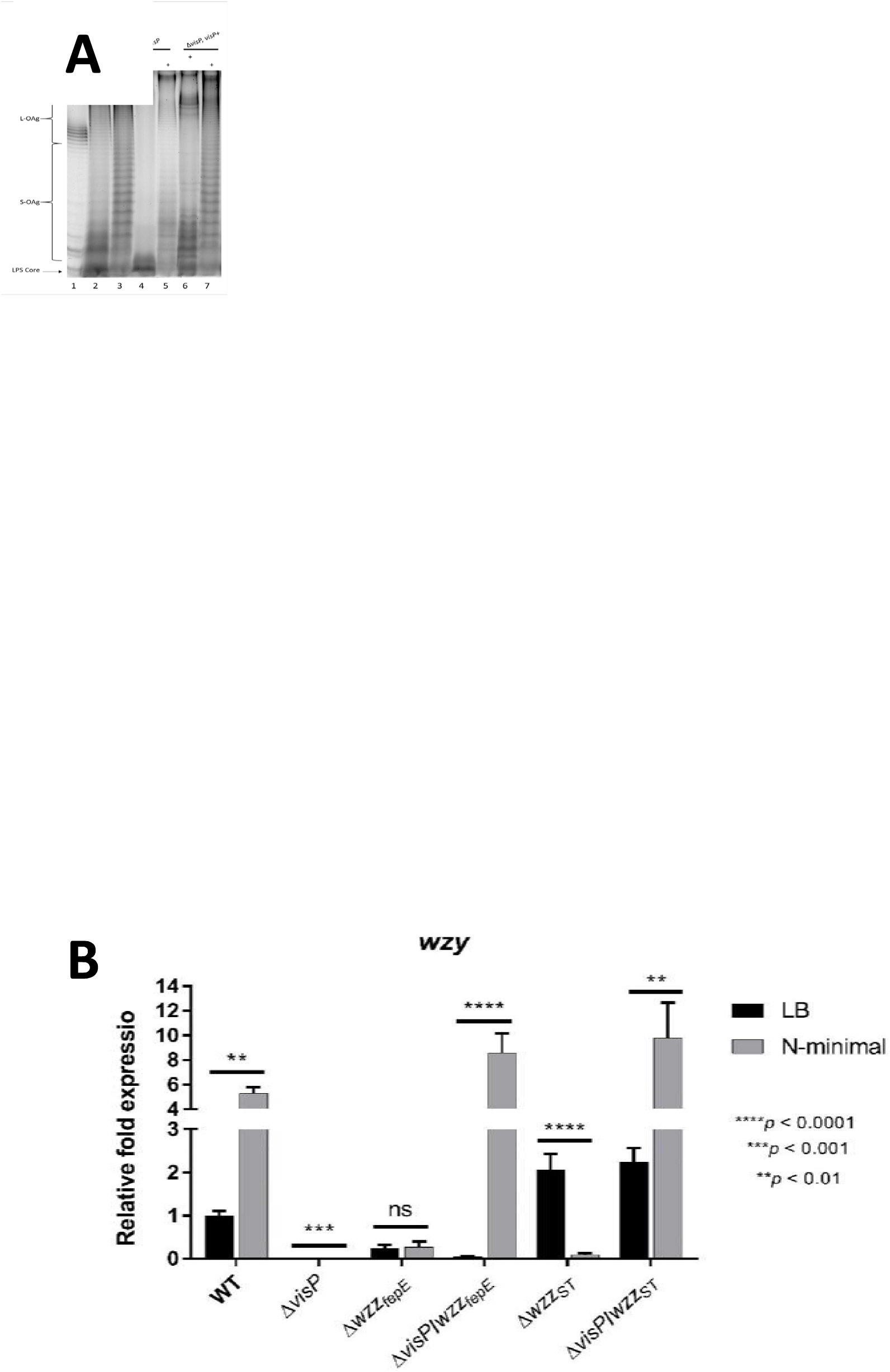
OAg changes are driven by intracellular conditions. (A) Electrophoresis profile of LPS in LB and N-minimal media. *E. coli* str. O55:B5 (lane 1), WT (lanes 2-3), Δ*visP* (lanes 4-5), and *visP*+ (lanes 6-7). (B) qRT-PCR analysis of *wzy* polymerase in LB and N-minimal media: relative fold expression in WT, Δ*visP*, Δ*wzz_fepE_*, Δ*wzz_ST_*, Δ*visP/wzz_fepE_* and Δ*visP/wzz_ST_* strains. ** *p* < 0.01 and **** *p* < 0.0001.

### Periplasmic VisP changes the OAg final structure

VisP is a periplasmic protein that is important for virulence and stress responses (61) and was initially described as a BOF family member (61, 64). OAg chain biosynthesis by the Wzy-dependent pathway mainly occurs in the periplasmic environment (52); based on this location, we assessed whether VisP plays a role in OAg chain formation. The Δ*visP* mutant strain exhibited a noticeable decrease in OAg chains in both growth media (Fig 1A, lanes 4 and 5; see also S1C and S1D Figs). In LB medium, this mutant appeared to have a single RU in LPS (Fig 1A, lanes 4; see also S1C Fig), a rough-like phenotype. The complemented strain *visP+* had a restored OAg profile that was similar to the WT profile and presented an increase in both VL- and S-forms in N-minimal medium; however, these levels were lower than those in the WT strain (Fig 1A, lanes 6 and 7; see also S1E and S1F Figs). Double mutants of *visP* and the PCP genes, *wzz_ST_* and *wzz_fepE_*, were designed to address the role of VisP as a protein important for the assembly of OAg at the final length observed here (Fig 1A) and to investigate the extremely low *wzy* expression levels in the single Δ*visP* mutant (Fig 1B). Again, *wzy* expression levels were highly upregulated in N-minimal growth medium compared with those in LB medium for both double mutants (Δ*visP*/*wzz_fepE_* and Δ*visP*/*wzz_ST_*) and the WT strain (Fig 1B). However, in LB medium, *wzy* expression levels diverged between the double mutants; compared to that in the WT, *wzy* expression was downregulated in Δ*visP/*Δ*wzz_fepE_* (similar to the Δ*visP* single mutant) and upregulated in Δ*visP/*Δ*wzz_ST_* (Fig 1B). Hence, the extreme downregulation of *wzy* in the absence of VisP was reversed by deleting one of the PCP genes with a major response to *wzz_ST_* deletion under nutrient-rich growth conditions. Next, we further evaluated macrophage-bacteria interactions during different conditions to explore the aspects of OAg assembly during intracellular environment survival.

### Changes in intracellular stress conditions and the role of PCP in pathogenesis

The *pmrB* gene, also known as *basS*, encodes a two-component system (TCS) sensor kinase that transcriptionally regulates chemical modifications of LPS upon sensing different environmental levels of Fe^3+^, Al^3+^ and acidic pH (73-75). Thus, *pmrB* plays a role in the control of the expression of the PCP genes *wzz_ST_* and *wzz_fepE_* (76). In *S*. Typhimurium, the transcriptional regulation of the *wzy* gene has not been elucidated. The *phoP* gene encodes a global TCS sensor kinase that regulates bacterial LPS remodeling through interplay with PmrAB (77). This TCS also helps regulate metal uptake (40, 78, 79), SPI-2 virulence effector expression (46) and resistance to cationic antimicrobial peptides (80, 81). All genes cited above exhibited undetectable expression levels in the Δ*visP* mutant background, in contrast to WT (Fig 2A). Complementation with *visP* partially restored the expression of *pmrB*, the two PCPs and *phoP* (Fig 2A). Therefore, VisP has a transcriptional effect on the genes of the Wzy-dependent OAg chain biosynthesis pathway, which is reflected in the OAg profile (Fig 1A, lane 4). As the absence of VisP results in modifications of LPS, the importance of VisP was then evaluated during intracellular survival within J774 macrophages. We performed distinct macrophage cell survival assays (67, 82, 83) to analyze the different stages of bacteria-macrophage interactions during the infection process. First, we performed a phagocytosis assay to quantify the bacterial load directly engulfed by J774 macrophages without extra time after bacteria-macrophage interactions and associated the load with the OAg pattern observed in the Δ*visP* mutant (Fig 1A, lanes 4 and 5). The absence of longer OAg forms in the Δ*visP* mutant resulted in an increase in macrophage bacterial uptake of more than one order of magnitude compared to WT levels (Fig 2B). A similar result was observed for the OAg-absent Δ*waaL* mutant, which had been previously reported (35) as evidence of the role of OAg chains in macrophage evasion. Complementation with the *visP* gene restored the phenotype to WT levels (Fig 2B). Then, we evaluated macrophage uptake and survival during a 3-h intracellular replication assay. In contrast to the previous assay, the Δ*visP* single mutant showed a statistically significant decrease in J774 macrophage internalization of 1.5 orders of magnitude compared to that in the WT (Fig 2C), as previously reported (61). *visP* gene complementation restored the WT phenotype (Fig 2C). An extended 16-h intramacrophage replication assay was subsequently performed (Fig 2D) to evaluate OM landscape remodeling in the SCV environment, which is similar to N-minimal growth conditions (Fig 1A). The semirough OAg Δ*visP* mutant strain exhibited a decrease of a half-order of magnitude compared to the WT strain (Fig 2D), a much smaller difference than that in the previous assay (Fig 2C). Δ*visP* mutant complementation restored the WT intramacrophage replication phenotype (Fig 2D). After observing these different phenotypes, gene expression levels from specific targets associated with intramacrophage survival and replication were evaluated to investigate whether the observed differences were direct effects of OAg and other SCV environmental features. Initially, we assessed the expression levels of *sifA*, an important SPI-2 TTSS effector related to SCV maintenance and the formation of cellular *Salmonella*-inducing filaments (84), under N-minimal bacterial growth conditions. The Δ*visP* mutant presented extremely low levels of *sifA* expression compared with the WT levels (Fig 2E). Expression levels similar to WT were restored by *visP* complementation (Fig 2E). The SPI-3 gene *mgtB* encodes an Mg^2+^ transporter ATPase related to intracellular adaptation and survival (85, 86). We observed very low *mgtB* expression levels in the Δ*visP* mutant under N-minimal growth conditions compared to the WT levels (Fig 2F). The complemented strain *visP*+ partially restored *mgtB* expression to WT levels (Fig 2F). The transcriptional downregulation of the genes *sifA* and *mgtB* was in accordance with the reduced 3-h intracellular replication phenotype of the Δ*visP* mutant, indicating a clear decrease in *S*. Typhimurium virulence. Furthermore, the absence of the OAg chain reduced the protection for this strain against macrophage-mediated phagocytosis. However, the Δ*visP* mutant was able to remodel its OM under conditions resembling the SCV environment, producing longer OAg forms; however, this production did not reach WT levels (Fig 1A, lane 5; and S1D Fig). The phenotypic difference between the Δ*visP* and WT strains decreased after 16 h of intracellular replication when they were compared in a 3-h assay, even though the Δ*visP* mutant downregulated intramacrophage survival-related genes. This late adaptation of the Δ*visP* strain may be related to longer chain OAg recovery under growth conditions resembling the SCV environment, which may confer greater protection against macrophage-mediated phagocytosis.

**Fig 2.**
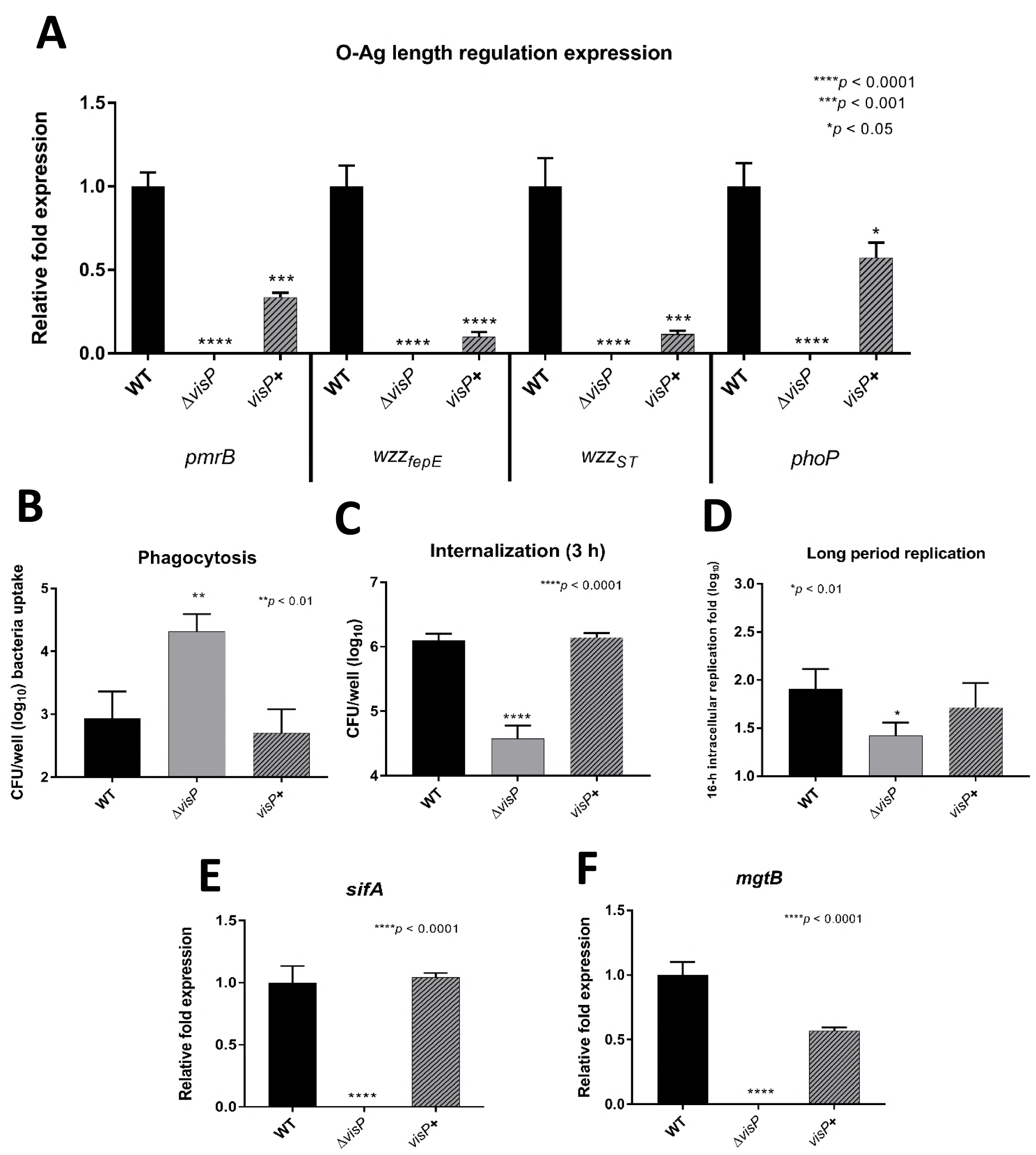
Roles of OAg and VisP in *S*. Typhimurium pathogenesis. (A) qRT-PCR analysis of genes involved in the OAg biosynthesis pathway: relative fold expression of *pmrB*, *wzz_fepE_*, *wzz_ST_*, and *phoP* genes in WT, Δ*visP* and *visP*+ strains grown in LB. (B) J774 phagocytosis, (C) internalization and (D) long-period replication of WT, Δ*visP* and *visP*+ strains. qRT-PCR analysis of (E) *sifA* (N-minimal growth condition) and (F) *mgtB* (LB growth condition) genes: relative fold expression in WT, Δ*visP* and *visP*+ strains. * *p* < 0.05, ** *p* < 0.01, *** *p* < 0.001, and **** *p* < 0.0001.

### The role of VisP-dependent PCP in the final OAg structure

The PCPs, together with Wzy polymerase, are responsible for finalizing OAg chain assembly. Initially, we addressed the two PCP single mutants, Δ*wzz_ST_* and Δ*wzz_fepE_*, during *wzy* gene expression analysis to better understand the role of VisP and the PCPs in this process under nutrient-rich growth conditions (LB medium). Compared to the results in the WT strain, the change in *wzy* expression in the Δ*visP/*Δ*wzz_ST_* was similar to those in the respective PCP single mutants, in contrast to Δ*visP/wzz_fepE_*, which exhibited lower *wzy* expression than Δ*wzz_fepE_* (Fig 1B). VisP and the PCPs participate in *wzy* transcriptional regulation; however, in the case of Wzz_ST_, the effect of PCP appeared to supersede that of VisP under LB growth conditions. Environmental conditions influenced the effects of VisP and PCP on *wzy* transcription, since under N-minimal growth conditions, *wzy* overexpression occurred only in the absence of both VisP and any OAg PCP, resembling WT levels (Fig 1B). Moreover, these three proteins appear to generate a balance in which VisP, Wzz_ST_, and Wzz_fepE_ are important during *wzy* expression. PCP gene expression levels in both double mutants compared to WT under different growth conditions were evaluated to explore the transcriptional regulation of the remaining PCP. Δ*visP/wzz_fepE_* exhibited a decrease in *wzz_ST_* levels in LB medium, whereas in N-minimal medium, its expression increased compared to that in the WT strain (Fig 3A). In contrast, Δ*visP/*Δ*wzz_ST_* displayed *wzz_fepE_* expression levels similar to those observed in the WT strain in LB medium, whereas the *wzz_fepE_* expression level greatly decreased in N-minimal medium (Fig 3A). Differentiated intestinal conditions also play an essential role in the absence of VisP, providing partial complementation of the VL- and S-forms in the Δ*visP/*Δ*wzz_ST_* mutant under N-minimal growth conditions (Fig 3B). Transcriptionally, N-minimal medium appeared to favor the Wzz_ST_ PCP, whereas *wzz_fepE_* expression was decreased in this medium. LB medium favored Wzz_fepE_, whereas *wzz_ST_* was more highly expressed in N-minimal growth conditions, but this expression always occurred in the absence of VisP. However, the VL- and S-forms were clearly phenotypically distinguishable under N-minimal conditions, which may be correlated with *wzy* overexpression (Fig 1B). Together, these results indicate complementary functions of the two PCPs based on distinct surroundings and nutrient conditions.

**Fig 3.**
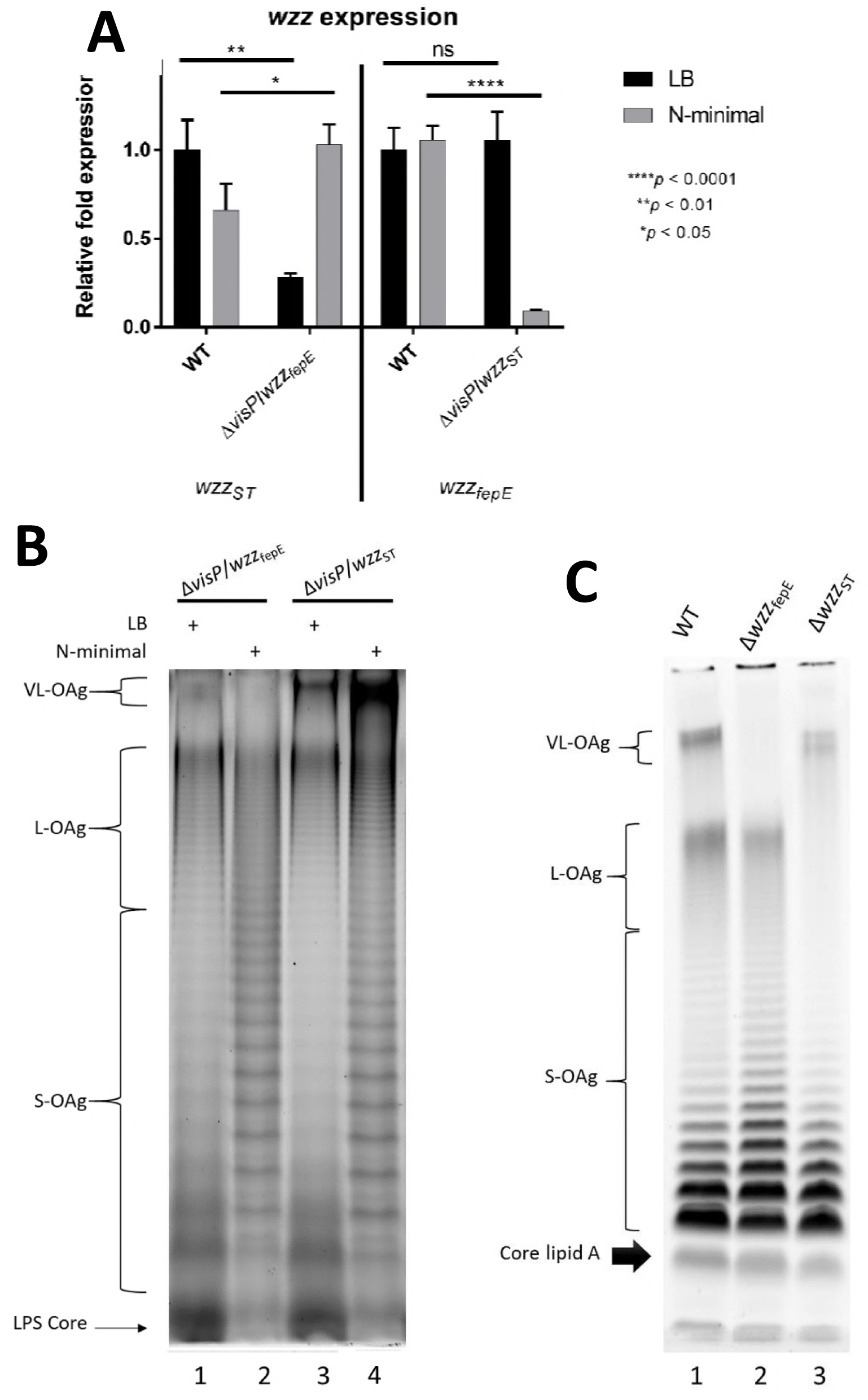
The interplay of VisP and PCP determines the final length of OAg. qRT-PCR analysis of (A) *wzz* expression in LB and N-minimal media: relative fold expression in WT, Δ*visP/wzz_fepE_* and Δ*visP/wzz_ST_* strains. (B) High-resolution electrophoresis (15% SDS-PAGE) and staining with a ProQ Emerald 300 LPS gel staining kit (Thermo Fisher, Massachusetts, USA) of the LPS profiles of the following strains grown in LB and N-minimal media: Δ*visP/wzz_fepE_* (lanes 1-2) and Δ*visP/wzz_ST_* (lanes 3-4). (C) Gradient SDS-PAGE (4-12%) of the LPS profile stained with a ProQ Emerald 300 LPS gel staining kit (Thermo Fisher, Massachusetts, USA) showing the core lipid A and different OAg chain lengths in WT, Δ*wzz_fepE_* and Δ*wzz_ST_* strains, all grown in LB. * *p* < 0.05, ** *p* < 0.01, *** *p* < 0.001, and **** *p* < 0.0001.

### Novel OAg length control

The Δ*visP* mutant was previously reported to have differentiated lipid A modification mediated by the LpxO enzyme (87). The single mutant Δ*visP* also showed a clearly distinct OAg phenotype with different patterns depending on the growth condition (Fig 1A). The two double mutants of *visP* and one of the genes encoding a PCP protein showed differentiated OAg profiles compared with those of their respective single PCP gene mutants (Fig 3B and 3C). The Δ*visP/wzz_fepE_* strain did not show the VL-OAg banding pattern (Fig B, lanes 1 and 2), consistent with the Δ*wzz_fepE_* phenotype (Fig 3C, lane 2) and the absence of Wzz_fepE_. Surprisingly, Δ*visP/wzz_ST_* showed an unprecedented phenotype in both media (Fig 3B, lanes 3 and 4), with restored L-forms compared to that in the Δ*wzz_ST_* single mutant (Fig 3C, lane 3). This finding contradicts the established consensus on Wzz_ST_-dependent L-OAg chain formation (57-59); Wzz_fepE_ is the remaining PCP in this background and may exert this function by cross-complementation. Moreover, the S-forms were very noticeable between different growth conditions; specifically, the S-forms showed a well-defined profile in N-minimal medium (Fig 3B). Together, these results suggest that the novel role of Wzz_fepE_ seems to be observed only in the absence of periplasmic VisP.

### Longer OAg facilitates intestinal colonization

*S*. Typhimurium causes gastroenteritis and promotes colitis in murine models (88, 89). Therefore, we evaluated the effects of OAg modal length patterns in an *S*. Typhimurium colitis model via the oral route in streptomycin-pretreated C57BL/6 mice (90). First, the semirough OAg phenotype (91, 92), which was observed in the Δ*visP* mutant, was attenuated by more than two orders of magnitude during *in vivo* colonization compared with the WT levels, as noted both 1 and 2 days postinfection (p.i.) (Figs 4A and 4B). These results were similar to those previously described in a Balb/C mouse colitis infection model (61). In the *in vivo* assays, the Δ*visP*/*wzz_fepE_* double mutant showed an increase in colonization of more than 1 order of magnitude in comparison with WT levels at day 1 p.i. (Fig 4A). These data demonstrate that the VL-OAg-absent Δ*visP*/*wzz_fepE_* mutant strain was better able than WT to adapt to host defenses and intestinal colonization *in vivo* at day 1 p.i. (Fig 4A). However, this mutant showed a decrease in colonization of half of an order of magnitude at day 2 compared to the level at day 1 p.i. (Figs 4A and 4B). The Δ*visP*/*wzz_ST_* mutant presented a more prominent two-order-of-magnitude increase between day 1 and 2 p.i. during murine intestinal colonization (Figs 4A and 4B). This limited colonization may be due to the absence of VL-OAg in the Δ*visP*/*wzz_fepE_* mutant strain. The only exception was the Δ*visP*/*wzz_ST_* double mutant strain, which showed higher colonization at day 2 p.i. than day 1 p.i. (Figs 4A and 4B). In addition to intestinal colonization fitness, we measured the transcriptional levels of two distinct interleukins, IL-17A and IL-22, to better assess the induction of inflammation by the following strains: WT, the single mutants Δ*wzz_fepE_* and Δ*wzz_ST_*, and the double mutants Δ*visP*/*wzz_fepE_* and Δ*visP*/*wzz_ST_* at two days p.i. in the colon (Fig 4C and 4D). IL-17A mediates neutrophil recruitment (93, 94) and suppresses *S*. Typhimurium colonization in enteric mucosa by inducing the production of antimicrobial peptides at the epithelial surface (95). The transcriptional levels of *Il17a* in the Δ*wzz_fepE_* and Δ*visP*/*wzz_ST_* strains were similar to those in WT, whereas Δ*wzz_ST_* infection resulted in a twofold increase (Fig 4C). The double mutant Δ*visP*/*wzz_fepE_* exhibited a twofold reduction in *Il17a* expression in the colon compared to that in the WT strain (Fig 4C), which may be related to the success of the initial intestinal colonization at day one p.i. (Fig 4A). IL-22 is responsible for inducing the production of antimicrobial proteins in the mucosa, which sequester metal ions required for the mechanism of evasion by *S*. Typhimurium, allowing this pathogen to outcompete the microbiota, enhancing its colonization (96). Interestingly, both VL-OAg-absent strains, Δ*wzz_fepE_* and Δ*visP*/*wzz_fepE_*, induced less *Il22* expression than WT (Fig 4D), whereas the Δ*wzz_ST_* and Δ*visP*/*wzz_ST_* strains exhibited higher induction of *Il22* expression than WT (Fig 4D). During *Salmonella*-induced colitis, VL-OAg also plays a role in bile salts resistance (97). Accordingly, we performed a bile resistance assay (98) comparing previous LB with N-minimal growth. The WT and Δ*visP* backgrounds showed growth recovery after 40 min at a lethal bile salt concentration (S2A and S2B Fig) only after previous N-minimal growth, which enhances VL-OAg chains (Fig 1A). After one hour at a lethal bile salt concentration, the longer OAg chain-deficient strain Δ*visP* exhibited a two-order-of-magnitude decrease in CFUs compared with that in the WT (S2C Fig). The low bile salts resistance may diminish the bacterial load of Δ*visP* reaching the intestines and affect the day one p.i. intestinal colonization levels (Fig 4A). The higher levels of IL-22 production and the trimodal OAg pattern similar to that of WT (Figs 1A and 3C.1) may explain the Δ*visP*/*wzz_ST_* strain colonization fitness at two days p.i. (Fig 4B), and the performance of this strain here confirms the importance of the L- and VL-OAg forms during the *in vivo* infectious process.

**Fig 4.**
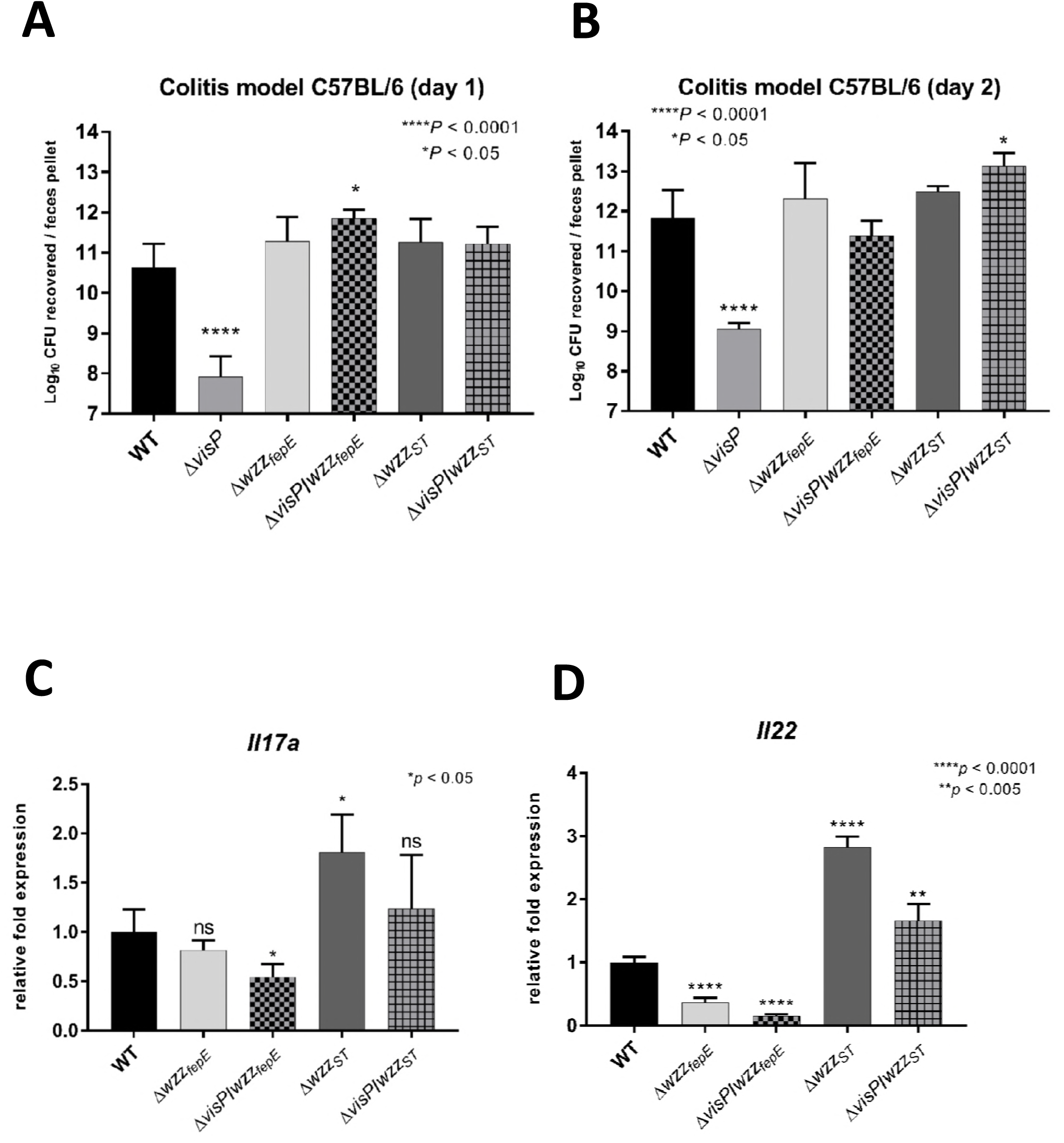
Longer OAg facilitates intestinal colonization. Colitis model of infection performed in C57BL/6 mice. (A) Day 1 p.i: log10 CFU of the WT, Δ*visP*, Δ*wzz_fepE_*, Δ*wzz_ST_* Δ*visP/wzz_fepE_* and Δ*visP/wzz_ST_* strains recovered from each feces pellet. (B) Day 2 p.i: log10 CFU of the WT, Δ*visP*, Δ*wzz_fepE_*, Δ*wzz_ST_* Δ*visP/wzz_fepE_* and Δ*visP/wzz_ST_* strains recovered from each feces pellet. qRT-PCR analysis of *Il17a* (C) and *Il22* (D) performed with RNA extracted from the mouse colon: relative fold expression in WT, Δ*wzz_fepE_,* Δ*wzz_ST_*, Δ*visP/wzz_fepE_* and Δ*visP/wzz_ST_* strains causing colitis. * *p* < 0.05, ** *p* < 0.01 and **** *p* < 0.0001.

### Flagellar changes with different OAg

Flagella play an important role in *S*. Typhimurium pathogenesis by helping the bacteria move toward nutrient-rich environments and eliciting inflammation processes in the gut (5, 6, 8-10). The flagella regulon is formed by 25 operons (over 67 genes) divided into 3 regulatory classes and is highly controlled by external factors. The class I operon *flhDC* produces FlhD and FlhC, components of the heteromultimeric complex FlhD_4_C_2_, which activates class II transcription through σ^70^ and autorepresses *flhDC* transcription. Hook basal body assembly is dependent on class II genes. After its completion, late substrates, such as FlgM, are exported from the cell, and σ^28^ starts the transcription of class III promoters. Class III genes include structural filament genes, such as *fliC* and *fljB*, and genes from the chemosensory pathway (99). Flagella gene expression was employed to better evaluate the full role of flagella structure and regulation via qRT-PCR of flagella essential genes from this complex regulon, such as the structural flagellin gene *fliC* (99), the master regulator *flhDC* (100, 101), genes such as *fliA* (102), which are responsible for class 3 transcription, and the gene encoding flagella proton motive force rotation, *motA* (103, 104). The Δ*visP* mutant showed very low levels of all tested flagellar genes (Figs 5A and 5B), and the complemented strain *visP*+ restored WT expression levels. We performed a swimming assay in semisolid agar to assess the motility of the strains and evaluate bacterial movement *in vitro*; after 8-h assays, five strains presented phenotypes similar to WT (Fig 5B). The Δ*visP* single mutant showed decreased motility compared to the WT levels. The complemented strain *visP*+ exhibited restored motility (Fig 5B). The *Salmonella* flagellin FliC, which constitutes the flagella filament subunit, was evaluated in terms of FliC protein levels via an immunoblotting assay with an anti-FliC monoclonal antibody and *fliC* gene expression levels via qRT-PCR. The FliC protein was not detected in Δ*visP*. Δ*visP/wzz_fepE_* presented a slightly reduced amount of flagellin relative to WT levels, and Δ*visP/wzz_ST_* overproduced flagellin FliC at abundant levels (Fig 5B). The protein levels matched *fliC* gene expression levels in all strains (Fig 5B). Considering the OAg forms, the semirough OAg strain Δ*visP* appeared to present less functional flagella, whereas the strains with more prominent VL-OAg, such as Δ*visP/wzz_ST_*, which have excess flagellin, did not seem to employ this abundant flagellin in a more efficient way than the WT strain (Fig 5B).

**Fig 5.**
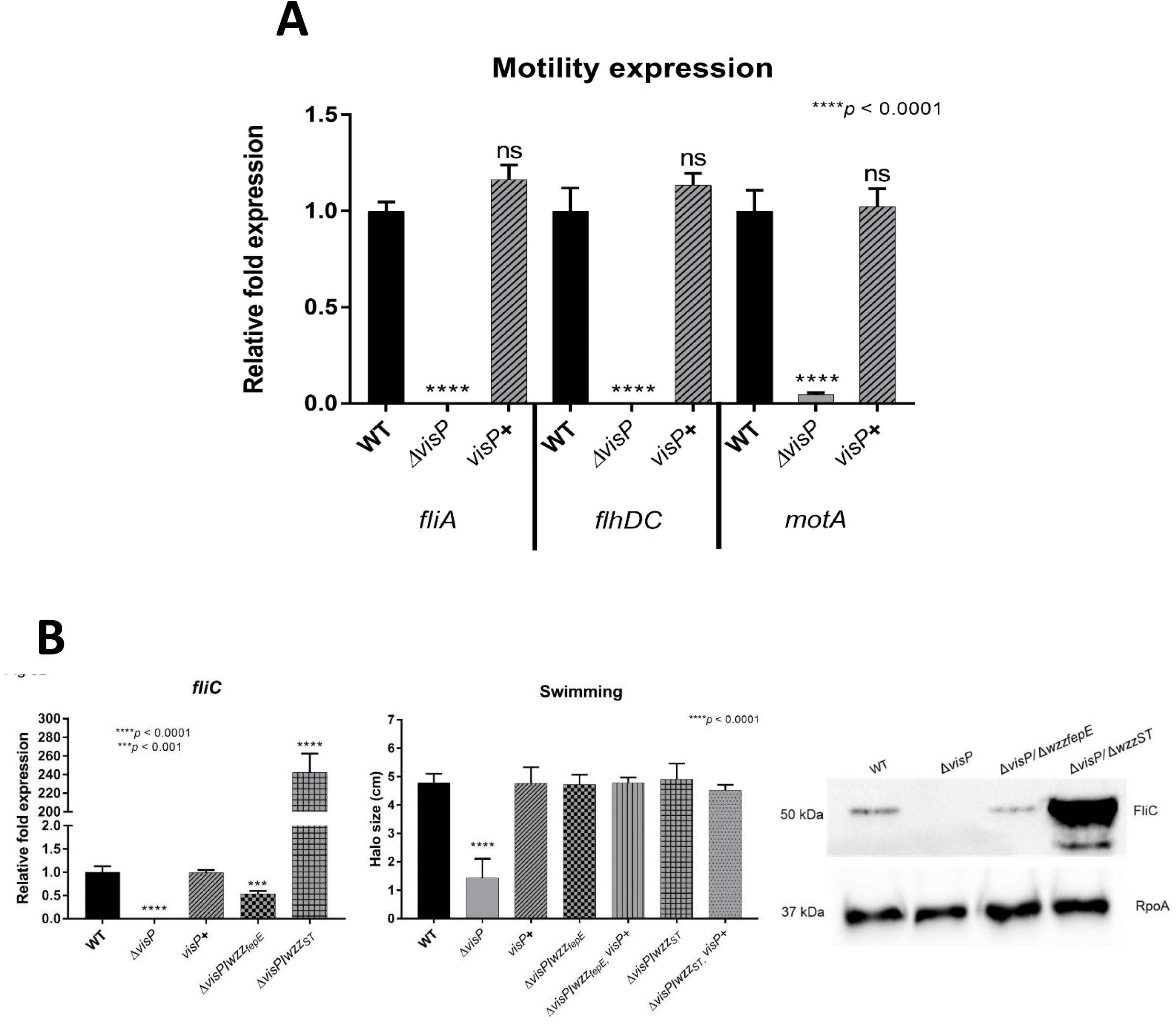
Changes in flagellar gene expression with different OAg. (A) qRT-PCR analysis of genes involved in flagellar assembly and motility: relative fold expression of the *fliA*, *flhDC* and *motA* genes in the WT, Δ*visP* and *visP*+ strains grown in LB. (B) qRT-PCR of the flagellin gene *fliC*, swimming motility assay and immunoblot performed with anti-FliC and anti-RpoA (control) monoclonal antibodies. *** *p* < 0.001 and **** *p* < 0.0001.

## Discussion

*S*. Typhimurium pathogenesis within the host depends on a complex assortment of factors mediated by stimulus recognition, such as bacterial movement across the lumen and mechanisms of evasion of host immune responses, to ultimately establish intestinal colonization. *S*. Typhimurium can elicit inflammation and benefits from this process. The host response through the innate immune system, among other defense machineries, recruits macrophage cells to combat the pathogen. Thus, *S*. Typhimurium has vast resources for colonization, as do other very successful bacterial pathogens, which employ several different virulence traits as an arsenal to strike back. The bacterial membrane plays an essential role in this process, providing protection against host defense mechanisms and physical stability to multiprotomeric complexes to support bacterial homeostasis and sensing of surrounding stimuli, which all culminate in metabolic and structural changes.

Here, we have demonstrated the outstanding mechanism by which *S*. Typhimurium differentially shapes its OM landscape during conditions that resemble the intracellular environment. Clearly, this mechanism involves modifications of distinct OAg chain patterns, with increases in the VL-OAg (>100 RUs) and S-OAg (less than 16 RUs) forms (Fig 1A) of LPS. These environmental stimuli-mediated LPS changes were observed by Lateenmaki and colleagues (105), who also measured OAg profiles extracted from macrophage-bacteria cocultures with more evident VL-OAg augmentation and, conversely, L-OAg reduction. Other growth conditions, such as inactivated guinea pig sera and iron-limited media, also augment VL-OAg chains in *S*. Typhimurium, although the specific stimuli were not identified (31). The length of longer OAg forms is controlled by the PCP proteins Wzz_ST_ and Wzz_fepE_ in a possible interaction with Wzy polymerase, as previously described in *Shigella flexneri* (106). However, the underlying mechanism remains unclear. The distribution of OAg polymerization catalysis occurs via Wzy in a “catch-and-release” process in which Wzy adds each OAg RU or oligomer to a single RU chain end without remaining bound to the growing chain (53, 54, 56). This process results in a random OAg chain length distribution controlled by the OAg RU availability surrounding Wzy; therefore, short OAg forms should be most prevalent. The PCP Wzz_ST_ is the molecular ruler responsible for L-OAg forms, shifting the random distribution of OAg chains by approximately 30 RUs (58). Wzz_fepE_ is a second PCP molecular ruler that shifts the OAg distribution to chain lengths larger than 100 RUs (S1 Fig), as previously described (60). N-minimal medium is commonly employed for the expression of intracellular virulence traits, such as SPI-2 TTSS effectors (65), because it resembles the SCV environment. Here, we observed that *wzy* overexpression is mediated by N-minimal condition stimuli (Fig 1B). Taken together, longer OAg chains result from the increase in Wzy that arises from the OAg polymer elongation rate, which is due to its distributive mechanism of action (54) as predicted by Michaelis-Menten enzyme kinetics. Therefore, WT *S*. Typhimurium increases VL-OAg by upregulating *wzy*, as observed previously in *S. flexneri* (107), which increases OAg chain elongation; however, it maintains Wzz_fepE_ levels and reduces Wzz_ST_ levels, as PCP proteins may act as competitors in Wzy interactions.

Based on the OAg profile, we identified a potential novel role of periplasmic VisP as a missing piece in the Wzy-dependent OAg biosynthesis pathway puzzle. These findings provide novel insights into the complex mechanism of defined L-OAg and VL-OAg formation in *S.* Typhimurium. The absence of VisP induced a semirough OAg phenotype in LB nutrient-rich growth (Fig 1A, lane 4), which was caused by Wzy and PCP deficiency (Fig 1B). However, the Wzx flippase and WaaL OAg ligase do not appear to be affected, nor is the OM LPS export apparatus, as evidenced by the presence of LPS with at least a single OAg RU, such as in the S-OAg form (Fig 1A, lane 4). The absence of VisP appears to broadly affect *S*. Typhimurium transcription, based on the observed downregulation of OAg biosynthesis pathway genes, the SPI-2 TTSS effector gene *sifA*, SPI-3 *mgtB*, and flagella regulation (Figs 1B, 2A, 2E, 2F, 5A and 5B). This transcriptional modulation could be associated with a possible signal transduction pathway mediated indirectly by VisP, as this periplasmic protein is regulated by the QseC sensor kinase and has a cryptic Ara-C-like regulator encoded on the same operon as YgiV (61). Two other very important TCSs of *S.* Typhimurium are implicated, namely, PhoPQ and PmrAB, which are represented here by the expression of *phoP* and *pmrB*; these gene levels were equally downregulated in the Δ*visP* mutant (Fig 2A). Previously, these two TCSs were not considered important for sensor kinase QseC regulation (44). Our results indicate that TCS may exhibit cross-talk under specific conditions that are dependent on the membrane functions examined here, such as OAg chain length determination. Moreover, the probable VisP signal transduction effect is associated with PCP proteins and is exemplified by the transcriptional regulation of *wzy* in the Δ*visP* single mutant compared with that in the Δ*visP/wzz_fepE_* and Δ*visP/wzz_ST_* double mutants (Fig 1B). A recent structural study of the *E. coli* PCP proteins FepE and WzzE and *Salmonella enterica* Wzz_ST_ characterized a conserved G-rich motif on the cytoplasmic part of the second transmembrane domain (TM2) as a putative tyrosine kinase (pfam13807) (108). The N-terminal cytoplasmic region immediately before the transmembrane helix is highly conserved among PCPs and could be a potential domain for interactions with cytoplasmic proteins and signal transduction pathways (108). Current evidence suggests that VisP and both PCPs, Wzz_ST_ and Wzz_fepE_, work together during Wzy expression, which is the major control factor for OAg length. Double mutation of *visP* with *wzz_ST_* or *wzz_fepE_* resulted in increased *wzy* expression compared with that in the OAg semirough Δ*visP* mutant. Unexpectedly, L-OAg formation was Wzz_ST_-independent in the Δ*visP/wzz_ST_* double mutant (Fig 3B; see also Fig S1I and S1J), in contrast to expectations for OAg length determinants based on the previously described Wzz_ST_-dependent L-OAg biosynthetic pathway (57-59). Based on this finding, we presume that Wzz_fepE_ has a novel role during L-OAg biosynthesis in the absence of VisP via cross-complementation. Recently, enterobacterial common antigen (ECA) was first characterized in *Shigella sonnei* as linked to LPS (more specifically, to lipid A) anchored to the OM (109). Therefore, the L-OAg modality observed here in the absence of Wzz_ST_ in the Δ*visP/wzz_ST_* strain may be ECA_LPS_ chains with similar lengths of L-OAg. The hypothesis presented here is supported by the similar molecular weights of ECA and OAg RU, approximately 607 and 606 Da, respectively. Similar to the OAg system of *S*. Typhimurium, the system for ECA binding to LPS has its own polymerase (WzyE) and PCP protein (WzzE), which have been characterized only by genomic studies (110). Together, these findings indicate bacterial membrane heterogeneity and the simultaneous formation of glycolipid structures in the periplasm, a transient and rich environment with many distinct proteins that may interact with one another and participate in similar synthesis processes.

The Δ*visP* single mutant displayed a clear deficiency during mouse gut colonization in our colitis infectious model (Fig 4). The absence of VisP causes strong acidic pH susceptibility, as previous *in vitro* tests have shown that its absence in Δ*visP* affects survival under acidic conditions (61). In *E. coli*, longer OAg forms maintain membrane stability and low pH tolerance (111). Furthermore, the Δ*visP* strain is defective in bile salts resistance (Fig S2C), in which OAg longer forms, primarily VL-OAg, have an important role (97). Thus, the semirough OAg phenotype of the Δ*visP* strain may increase its susceptibility to the low pH of the stomach, as well as bile salts, reducing the bacterial load that reaches the intestine. Moreover, a phenotype with reduced levels of highly immunogenic proteins, such as FliC flagellin, may eliminate the ability of the Δ*visP* strain to elicit inflammation processes, thus affecting its initial colonization ability (5, 6, 8-16, 19-22, 29, 112).

As described by Barthel and colleagues, the induction of the inflammation process by *S*. Typhimurium in a streptomycin-pretreated mouse colitis model was observed starting at 8 h p.i. based on the presence of PMN in the mouse intestinal lumen (90). Thereafter, they observed a large increase in PMN cell transmigration to the intestinal lumen from 20 h to 48 h p.i. (90) as a clear consequence of the inflammation process (5, 6, 8-16, 19-22, 29, 112). Therefore, a few h p.i., the host started to attack the pathogen through cells of the innate immune system. OAg chains function indispensably to trick macrophage recognition and consequent phagocytosis (Fig 2B). The OAg rough phenotype is caused by complete OAg absence in the OM as a result of impairment of Wzx flippase or WaaL ligase function (35, 113-118). The semirough OAg phenotype is characterized by LPS with only one OAg RU, which occurs due to Wzy malfunction (91). Both phenotypes result in a defective Gram-negative bacterial membrane, which enables more efficient antibody opsonization and activation of the complement cascade, culminating in increased macrophage-mediated phagocytosis (29, 30, 35, 55, 91, 119), as observed here in the Δ*visP* strain (Fig 2B).

*S*. Typhimurium must adapt itself to sense phagosome differences in acidification and nutrient starvation after macrophage-mediated phagocytosis, such as low Mg^2+^ concentrations, through TCSs such as PmrAB and PhoPQ. Together, these TCSs activate cascade responses to defend against host enzymes and counterattack by coating the SCVs (28, 40, 77-79, 81, 82, 120, 121). The modulation of these responses triggers SPI-2 T3SS, and its gene expression is essential because it encodes effector proteins that are injected into host cells to modulate and manipulate the cell machinery for the bacterium’s benefit (69-71, 84).

We showed that the long forms of OAg in *S*. Typhimurium play important roles within macrophages (Fig 2). Initially, we demonstrated that *S*. Typhimurium OM remodeling is driven by SCV environments resembling stimuli conditions, resulting in increased OAg chains, primarily the VL-OAg pattern (Fig 1A). The Δ*visP* strain has a semirough OAg phenotype and is less protected within macrophages, as reflected in its intracellular survival levels (Fig 2C). However, the Δ*visP* mutant is able to produce OAg longer forms under conditions resembling SCVs, but this production occurs to a lesser extent than it does in the WT strain (Fig 1A). Although Δ*visP* presents low expression of important intramacrophage survival genes, such as *sifA* and *mgtB*, its OM is remodeled with OAg long forms in the SCV, conferring sufficient protection to reduce its phenotypic differences from the WT strain in a 16-h assay (Fig 2D). Moreover, SPI-2 TTSS is assembled over longer periods, and some of its structures are detected only after 5 h upon SPI-2 gene expression induction (122). *S*. Typhimurium in intracellular macrophages may overproduce VL-OAg chains, which may help SPI-2 TTSS build and promote the stability of the SseBC appendage structure, a gigantic structure that is larger than 160 nm (122). Furthermore, SPI-2 TTSS is sheathed by protein effectors during and after its assembly, approaching SCV membranes and reaching the macrophage cytoplasm to exert its function in SCV maintenance (122). Consequently, VL-OAg chains may also help these effector proteins move across SPI-2 TTSS. An analogous function for VL-OAg was proposed in giant adhesin SiiE secretion, which pulls its structure through ionic forces mediated by Ca^2+^ cations present at the OAg chains (34). L-OAg chains employ an initial protective role and may provide more support for VL-OAg, maintaining LPS stability as a whole. More importantly, longer OAg forms confer strong protection against macrophage phagocytosis to *S*. Typhimurium, allowing the bacterium to survive in the phagosome and thus creating a propitious environment for its replication. Taking all these observations into consideration, we propose a model in which *S.* Typhimurium strengthens its protection by increasing OAg longer forms inside the phagosome through the overproduction of Wzy (Fig 6A). This process occurs via transcriptional control of *wzy*, which may be mediated by VisP and PCP protein interactions in the periplasm, creating a signal transduction cascade ignited by SCV environmental stimuli (Fig 6A).

**Fig 6.**
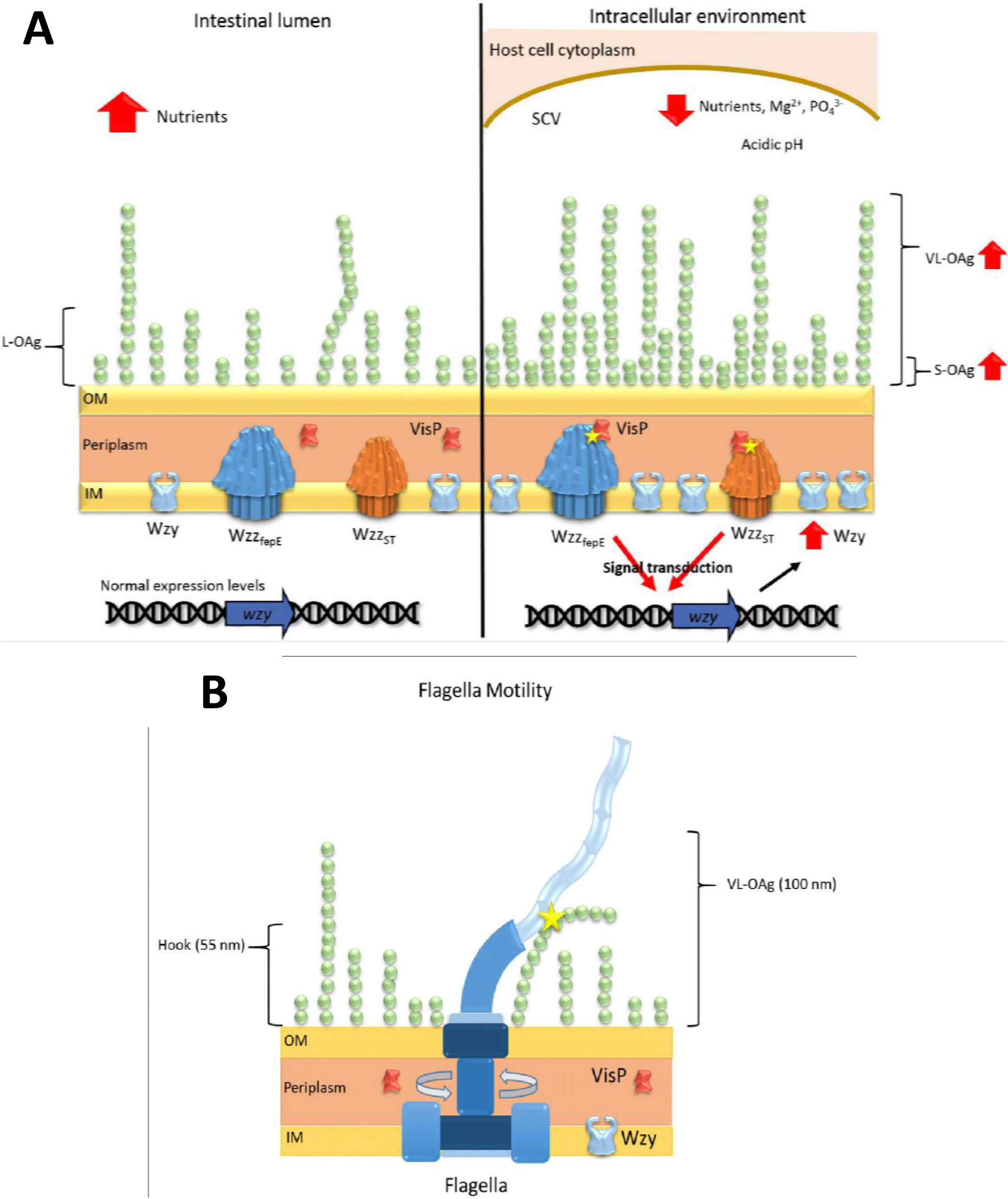
Proposed models of OAg outer-leaf remodeling and flagella/OAg interaction. Model of (A) OAg final chain length in the intestinal lumen (left) and intracellular environment (right). (B) *S*. Typhimurium flagella interaction with VL-OAg chains.

All PCP mutant strains caused similar *Il17a* expression levels in a mouse colitis model to those in the WT, with a slight reduction for *wzz_fepE_* mutants (Fig 4C). In this manner, we presume that the levels of intestinal inflammation and, more importantly, levels of neutrophil transmigration to the intestinal lumen are comparable for all strains. Nonetheless, distinct *Il22* transcription patterns were observed, with a significant decrease in mice infected by VL-OAg-absent strains (Fig 4D). IL22 induces the production of antimicrobial proteins responsible for metal ion starvation, such as lipocalin-2 (94). *S*. Typhimurium can overcome this host defense mechanism and thus outcompete the affected microbiota (96). Since infection with strains presenting VL-OAg resulted in greater *Il22* expression, this OAg modal length may induce the expression of *Il22*, culminating in more successful intestinal colonization. The higher expression rate of *Il22* combined with a more protected OM presenting a WT-like OAg profile (Fig 3B) could explain the increased level of intestinal colonization by Δ*visP*/*wzz_ST_* strain at 48 h p.i. (Fig 4B).

Bacterial motility may be considered one of the most complex assembly mechanisms in bacterial structures because it includes a regulon network of more than 50 distinct genes (99) that are involved and directly influenced by environmental differences in the expression of the flagellins FliC and FljB of *S.* Typhimurium. Here, we also observed a role of VisP during bacterial motility, as the absence of periplasmic VisP caused a direct defect during swimming. After gene complementation, the swimming ring size was restored to WT levels (Fig 5B). The Δ*visP* single mutant had very low expression levels of the master regulator *flhDC,* class 2 sigma factor *fliA,* class 3 gene *motA* and flagellin gene *fliC*. The FlhD_4_C_2_ heteromultimeric complex encoding *flhDC* expression starts the flagella assembly cascade, which is followed by class 2 and 3 flagella operons. The *fliA* gene encodes a sigma factor that activates the transcription of class 3 genes. After hook completion, FlgM is exported from the cell and allows σ^28^ to start class 3 transcription. The flagellin FliC is the main filament component and is also exported via flagellar T3SS (123). The MotAB system of flagella is responsible for the crucial motility of the flagella and is assembled via a Sec-dependent pathway (99). These two proteins form the ion-conducting stator complexes necessary for flagellar rotation (103, 104).

Mutation of either *wzz* gene in the Δ*visP* mutant background restored flagella gene expression to WT levels and the presence of the FliC protein in bacterial membrane extracts (Fig 5B). Previously, PCP proteins have been shown to have a dual regulatory role in motility networks in bacterial LPS modifications. The Wzz_ST_ protein is required to maintain the balance of 4-aminoarabinose and phosphoethanolamine in *Salmonella* lipid A modifications, which are mediated by ArnT and EptA, respectively (76). In *C. jejuni*, a gene encoding a phosphoethanolamine transferase was directly correlated with lipid A modifications and the flagellar rod protein FlgG (124).

The Δ*visP* single mutant lacked all flagella parameters evaluated here, such as the expression of various flagella genes and reduced production of FliC flagellin. These deficiencies may be directly linked to its attenuation in our *in vivo* and *in vitro* tests, such as in phagocytosis and short- and long-period macrophage survival. The Δ*visP/wzz_ST_* double mutant exhibited prominent overproduction of the flagellin FliC compared to WT levels; however, this overproduction apparently did not influence its motile phenotype (Fig 5B). The presence of FliC protein indicates mature flagella, and overproduction of the filament suggests that the double mutant strain Δ*visP/wzz_ST_* presents more flagella or longer filaments than WT, although this change may not be reflected as a more efficient swimming mechanism (Fig 5B). Thus, we propose that VL-OAg with sizes over 100 nm (60) physically interact with the rotating flagellum, as the hook structure is smaller than approximately 55 nm (125), to reach the maximum plateau of motility efficiency even in the presence of more flagella or longer filaments (Fig 6B). Previous studies have demonstrated the importance of LPS for surface or swarming motility (126), mainly due to the wettability of LPS. Moreover, the increase in expression levels of OAg related genes occurs concomitantly with the expression of class 3 motility genes and, consequently, flagella maturation (127). Wang *et al.* have also shown that the overexpression of the iron uptake system (127), encoded adjacent to the genome location of *wzz_fepE_* (60), is directly affected by different iron concentrations (31). Important LPS structural assembly genes, such as *rfaC*, impact the secretion of proteins in *S.* Typhimurium, such as SipA, FlgK, FliC, FliD, SipC, InvJ, and FlgL, thereby hampering protein translocation, which may be enhanced by the absence of OAg (128). In addition, proteins related to LPS modification also perform posttranslational modifications of flagella structural proteins, such as phosphoethanolamine addition to FlgG catalyzed by Cj0256 in *Campylobacter jejuni* (124) and flagellin O-glycosylation by the protein encoded by *rmlB* in *Burkholderia pseudomallei* (129). Similarly, we hypothesize that the Δ*visP/wzz_ST_* mutant strain may overproduce FliC flagellin in a correlated mechanism with PCP Wzz_fepE_ in the absence of both VisP and other PCP Wzz_ST_, supporting the importance of OAg chain structures during the secretion of other membrane proteins.

The periplasmic protein VisP appears to be essential for cell membrane maintenance, virulence, and proper flagella function. In addition, the Δ*visP/wzz_fepE_* double mutant was also able to partially restore flagellin FliC and *fliC* gene expression (Fig 5B). Together, these data show that VisP has a novel and important role during flagella expression that warrants further study and that both Wzz PCPs presented here may function as cofactors to restore flagellar function in *S*. Typhimurium via FliC. The VL-forms (Δ*visP/wzz_ST_*) apparently increase the overproduction of FliC flagellin, which may be correlated with the size of the LPS chains. Moreover, the L-forms (Δ*visP/wzz_fepE_*) induce greater production of FliC than in the Δ*visP* single mutant; therefore, there is a link between the longer forms of OAg and flagella production that is not fully understood and requires clarification.

The bacterial membrane is the first line of interaction with the host. Determining how it senses and responds to host cell signals by changing its structure and composition is the initial step toward fully understanding this complex relationship. A better understanding of these processes will shed light on how hosts and pathogens have coevolved and how we can develop new therapies to dismantle pathogen defenses. In this work, we identified a *S*. Typhimurium response mechanism against macrophage phagocytosis. The next step will be to elucidate how each piece of this complex process exerts its function, which will enable the design of target-specific drugs to block their activity.

## Materials and methods

### Ethics statement

All procedures were performed under supervision of a veterinarian with the approval of the Animal Ethics Committee of the School of Pharmaceutical Sciences (CEUA/FCF/CAr n° 23/2016). The study was conducted in accordance with the Brazilian Law for Animal Research n° 11.794, October 8^th^ 2008, Decree n° 6.899, July 15^th^ 2009.

### Strains and plasmids

All strains and plasmids used in this study are listed in S1 Table. Recombinant DNA and molecular biology techniques were performed as previously described (130). The oligonucleotides used in this study are listed in S2 Table.

### Construction of isogenic mutants

Construction of isogenic nonpolar *S.* Typhimurium SL1344 *wzz_fepE_* and *wzz_ST_* single mutants and *visP*/*wzz_fepE_* and *visP*/*wzz_ST_* double knockout mutants was achieved by using λ Red mutagenesis (131). The *visP*/*wzz_fepE_* and *visP*/*wzz_ST_* double mutants were complemented with the *visP* gene cloned into the vector pBADMycHisA (*Kpn*I and *Eco*RI) (61) to generate strains 184 and 185.

### Macrophage infection assays

Intramacrophage phagocytosis assays were performed as previously described (35). In the survival assay, the cells were washed and incubated at 37°C and 5% CO2 for 3 h without antibiotics (67, 82, 83). After extracellular bacteria killing in the intramacrophage replication assay, the cells were washed and incubated at 37°C and 5% CO2 for 16 h with 15 μg/mL gentamicin (35).

### Motility assays

The swimming motility behaviors of *S.* Typhimurium were assessed in semisolid plates at 37°C for 8 h, as previously described (44, 132).

### Quantitative real-time RT-PCR

Quantitative RT-PCR was performed as previously described (133).

### Colitis model infection with *S.* Typhimurium

Mice (C57BL/6, 7 to 9 weeks old, female) were infected orally for the colitis model as previously described (90).

### Western blot assay

Culture supernatants were obtained from strains grown statically in LB medium at 37°C overnight. The medium was removed by centrifugation, and the cells were resuspended in 10 mL of PBS buffer plus 100 μL of phenylmethylsulfonyl fluoride (100 mM). Bacteria were lysed by sonication, pelleted by centrifugation (4,200 × g, 10 min, 4°C) and analyzed by SDS-PAGE. Proteins were separated by SDS-PAGE and electrophoretically transferred to nitrocellulose membranes via a Semi Dry Transfer Cell (Bio-Rad, Hercules-CA). Immunoblots were probed with monoclonal anti-flagellin FliC (InvivoGen, San Diego-CA) 1:1,000 and anti-RpoA (Santa Cruz Biotechnology, Dallas-TX) 1:5,000. Bound antibody was detected with horseradish peroxidase-conjugated secondary antibody anti-mouse IgG 1:2,500 (Promega, Fitchburg-WI), followed by enhanced chemiluminescence substrate (ECL, Promega, Fitchburg-WI) and detection in Chemidoc MP (Bio-Rad, Hercules-CA).

### Analysis of LPS profile

Samples were grown overnight aerobically on LB medium at 37°C. Cell cultures were diluted 1:100 and grown on LB or N-minimal media at 37°C and 200 rpm until an OD600 of 1.0 was reached. Then, 1 mL of cells was harvested and centrifuged at 15,000 × *g* for 5 min. The pellets were resuspended in lysis buffer (2% β-mercaptoethanol, 2% SDS and 10% glycerol in 0.1 M Tris-HCl, pH 6.8). The suspension was boiled for 10 min and then treated with 1.25 μL of proteinase K (20 mg/mL) overnight at 55°C. LPS was separated on 15% SDS polyacrylamide gels using a Laemmli buffer system at 25 mA and visualized by gel staining using the ProQ Emerald 300 LPS gel staining kit (Thermo Fisher Scientific, Waltham-MA).

### Bile salts resistance assay

Bacteria were grown in LB or N-minimal media until reaching exponential phase (OD_600_ 0.6) and then centrifuged at 4000 × *g* for 10 min. The pellets were resuspended in LB broth with 30% bile salts. OD_600_ readings were acquired every 20 min for 1 h, with the first measurement occurring after resuspension (98). To compare the two different strains, samples initially grown on LB broth were washed twice with PBS immediately after exposure to bile salts for 1 h. The samples were serially diluted and plated for CFU enumeration.

## Acknowledgments

We would like to thank all faculty members and technicians of the Biological Sciences Department from UNESP for assistance in setting up the Moreira laboratory and especially for sharing reagents and equipment for our initial work. In particular, we would like to thank Professors Sandro Valentini and Fernando Pavan for their assistance.

